# Membrane-enhanced repulsive interactions regulate protein diffusion in cell-size space

**DOI:** 10.1101/2025.04.22.650139

**Authors:** Hiroki Sakuta, Yuki Kanakubo, Sakura Takada, Naoya Yanagisawa, Tatsuro Oda, Koichi Mayumi, Koichiro Sadakane, Kei Fujiwara, Miho Yanagisawa

**Affiliations:** Komaba Institute for Science, Graduate School of Arts and Sciences, The University of Tokyo, Komaba 3-8-1, Meguro, Tokyo 153-8902, Japan; Center for Complex Systems Biology, Universal Biology Institute, The University of Tokyo, Komaba 3-8-1, Meguro, Tokyo 153-8902, Japan; Department of Biosciences and Informatics, Faculty of Science and Technology, Keio University, 3-14-1 Hiyoshi, Kohoku, Yokohama, Kanagawa, 223-8522 Japan; The Institute for Solid State Physics, The University of Tokyo, 5-1-5 Kashiwanoha, Kashiwa, Chiba 277-8581, Japan; Faculty of Life and Medical Sciences, Doshisha University, 1-3 Tatara Miyakodani, Kyotanabe, Kyoto 610-0394, Japan; Department of Physics, Graduate School of Science, The University of Tokyo, Hongo 7-3-1, Bunkyo, Tokyo 113-0033, Japan

**Keywords:** protein mobility, macromolecular crowding, confinement, reaction-diffusion wave

## Abstract

Intracellular molecular organization is often explained by attractive interactions driving clustering and phase separation. Although consideration of repulsive forces is essential in physics, their roles remain unclear in cellular contexts. Here, we demonstrated the fundamental role of repulsion in regulating protein diffusion within cell-size space. By analyzing negatively charged protein diffusion in bulk solutions and in cell-size spaces with membranes, we revealed that membrane-enhanced repulsion inhibited protein diffusion in cell-size spaces. This was due to the amplified electrostatic interactions among proteins because of the large membrane area-to-volume ratio. Notably, ATP, a cellular central energy source, further inhibited protein diffusion in cell-size spaces, whereas protein wave propagation on the membrane counteracted this inhibition. These findings suggest an active regulatory mechanism restoring molecular mobility by dynamically adjusting membrane-enhanced repulsive forces. Our study challenges the traditional emphasis on attractive interactions, highlighting repulsion as a critical tunable factor governing molecular transport and spatial organization in cells.

**Teaser:** Protein diffusion within crowded cell-size spaces is tuned by membrane-enhanced repulsion and its active regulation.

## 1. Introduction

Intracellular molecular organization, from nanoscale clusters to microscale condensates, is essential for proper functioning. Nucleoli formed via liquid–liquid phase separation (LLPS), grow over time to microscale structures, in which ribosomal RNA synthesis and processing occur (1). Ion channels exhibit proper structures and gating functions via multimerization (2). These diverse structures have traditionally been explained by attractive intermolecular interactions, resulting in the extensive qualitative analysis of attractive forces in various binding assays (3).

Attractive interactions bring biomolecules into close proximity, effectively minimizing the intermolecular distance. However, these factors do not solely account for the spatial organization of biomolecules in the cytoplasm. Not all biomolecules aggregate, with many remaining dispersed, implying the presence of repulsive forces counteracting attraction. Under physiological conditions (approximately pH 7), most biomolecular surfaces carry a net negative charge (4), resulting in electrostatic repulsions inhibiting uncontrolled aggregation. However, acidification reduces this repulsion and promotes LLPS, a phenomenon linked to various biological processes, including yeast dormancy, cancer progression, metabolism, and signal transduction (5). Therefore, tuning of electrostatic repulsion, rather than attraction, is important to regulate the intracellular organization. The role of repulsion on cell–cell interactions is reported previously (6), however, specific effects of repulsive forces on molecular behaviors in the cytoplasm remain unclear (7).

The strength of the electrostatic repulsion between biomolecules, such as proteins, is modulated by pH levels and salt concentrations, which affect electrostatic screening. Additionally, surfaces of lipid membranes, which are the basic structures of cell membranes, also carry a net negative charge under physiological conditions (8, 9). This similarity suggests that negatively charged proteins and membranes compete for the available cations in the solution. However, it remains unclear how the high membrane area-to-volume ratio alters the repulsive forces between proteins.

In this study, we investigated the roles of electrostatic repulsion in intracellular environments by encapsulating a negatively charged protein, bovine serum albumin (BSA) (10), in an artificial cell covered with a negatively charged lipid membrane and analyzed its molecular diffusion (11). Our experimental and simulation results revealed that membrane confinement in cell-size spaces significantly hindered BSA diffusion in crowded environments due to excessive protein–protein repulsion. This was because the competition between proteins and membranes for available cations enhanced protein–protein repulsion in cell-size spaces. Moreover, propagation of Min protein waves on the membrane counteracted this inhibition, effectively restoring protein mobility. Our findings suggest that intermolecular repulsion through membranes plays a key role in controlling molecular diffusion and that cells possess active repulsive regulatory mechanisms to optimize intracellular diffusion and spatial organization.

## 2. Results

### 2-1. BSA diffusion in cell-size droplets

We mimicked the cellular environment using cell-size droplets containing BSA solutions enclosed by a lipid membrane (Fig. 1a). A crowded cellular environment was characterized by (i) macromolecular crowding and (ii) cell-size membrane confinement.

**Figure 1.**
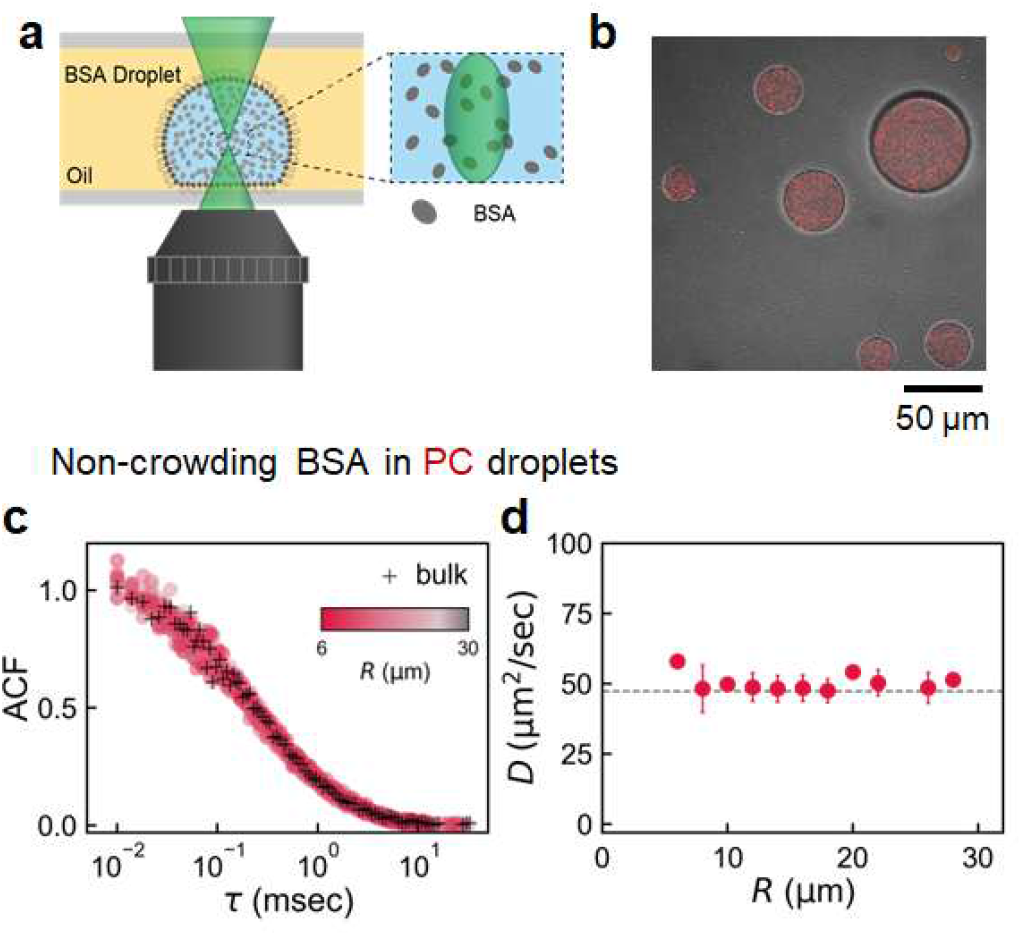
(a) Schematic illustration of a bovine serum albumin (BSA) droplet covered with a lipid membrane. The confocal region where the diffusion measurements are performed (enlarged image on the right) is in the center of the droplet at a height of 5 μm from the bottom. (b) Microscopic images of droplets containing 160 mg/mL BSA and 120 nM Texas Red (TR)-BSA (red). (c) Autocorrelation function (ACF) of TR-BSA diffusion in 10 mg/mL BSA in phosphate-buffered saline (PBS) is shown for bulk (black cross). (d) Various 1,2-dioleoyl-sn-glycero-3-phosphocholine (PC) droplets with radius (*R*) = 6–30 μm (red circle, left). Dependence of *D* (*n* = 25; error bars represent the standard deviation (S. D.)) on *R* was obtained by fitting for ACF (right). Average *D* value for droplets (49.3 ± 5.3 μm^2^/sec (Ave. ± S. D.; *n* = 25)) was almost similar to the bulk value indicated by the dashed line (*D*_bulk,_ = 47.3 ± 5.7 μm^2^/sec (Ave. ± S. D., *n* = 10)).

In our experiments, (i) macromolecular crowding was controlled by varying the BSA concentration and buffer conditions. Specifically, BSA concentrations of 10 and 160 mg/mL were used to represent the non-crowding and crowding conditions, respectively. A small amount of Texas Red-conjugated BSA (TR-BSA; 120 nM) was added to analyze protein diffusion via fluorescence correlation spectroscopy (FCS). In the absence of BSA, TR-BSA accumulated on the lipid membrane and glass surface. However, in the presence of 10 mg/mL BSA, TR-BSA was uniformly distributed within the droplets, as indicated via fluorescence microscopy (Fig. 1b). Therefore, 10 mg/mL BSA was used for the non-crowding conditions. A buffer with a pH of approximately 7 was used to prevent the aggregation or clustering of BSA due to electrostatic interactions. This condition was used to ensure that BSA maintained a negative net surface charge because its isoelectric point (pI) is approximately 4.7 (12). Two buffer types were employed to regulate the intermolecular repulsions: Phosphate-buffered saline (PBS) at pH 7.4 and 150 mM KCl, and 5 mM MgCl_2_ (hereafter referred to as the TKM) at pH 7.6, both with 150 mM NaCl and KCl (see Materials and Methods section 4-2).

For (ii) cell-size membrane confinement, droplet radius *R* and membrane lipid composition were varied. The lipid species included electrically neutral 1,2-dioleoyl-sn-glycero-3-phosphocholine (PC), negatively charged 1,2-dioleoyl-sn-glycero-3-phospho-(1’-rac-glycerol) (PG), and *Escherichia coli* extract polar (E. coli polar). Based on the negatively charged lipid contents, such as PG and cardiolipin contents, the membrane charge followed the order: PC (neutral lipids) > E. coli polar (containing 33% negative lipids) > PG.

First, we investigated the effect of (ii) cell-size membrane confinement on molecular diffusion without (i) macromolecular crowding. Figure 1c shows the representative Autocorrelation functions (ACFs) under 10 mg/mL BSA conditions in PBS in bulk systems (black) and within PC droplets with radius *R* of 6–30 μm (*n* = 25, red). The shade of red indicates the *R* value; the darker the shade, the smaller the *R* value. To minimize the effects of the high refractive index of the oil phase, measurements were conducted at the center of the partially adhered droplets at a height of 5 μm (Fig. 1a). ACF curves were consistent regardless of *R* values. To quantitatively analyze the diffusion coefficient *D* of TR-BSA, ACFs were fitted using Eq. (1) and derived *D* using Eq. (2). The resulting *D* values were plotted against *R* (Fig. 1d) and remained constant for different *R* values. Average *D* value in PC droplets was 49.3 ± 5.3 μm^2^/sec (Ave. ± S. D.; *n* = 25), consistent with the corresponding bulk values (dashed line; *D*_bulk,_ = 47.3 ± 5.7 μm^2^/sec (Ave. ± S. D.; *n* = 10). Similar *D* values in the bulk systems and droplets were obtained when TKM was used instead of PBS (Fig. S1; Table S1). These values align with the previously reported BSA diffusion coefficients of 53–60 μm^2^/sec (11, 13). Notably, *R*-independent *D* values of TR-BSA under non-crowding conditions suggest that (ii) cell-size membrane confinement has no significant impact on molecular diffusion under non-crowding conditions.

Next, we analyzed the synergistic effects of (i) macromolecular crowding and (ii) cell-size membrane confinement on molecular diffusion using droplets and bulk systems containing 160 mg/mL BSA in PBS. Figure 2a–f presents the ACFs and corresponding *D* values obtained by fitting the ACFs with Eq. 1 plotted against the droplet radius *R* (see section 4-6 in Materials and Methods). *D* value was normalized its corresponding bulk value (*D*/*D*_bulk_) (Fig. 2b, d, and f). Different colors represent the lipid types forming the droplet membranes: PC (red; panels a and b), E. coli polar (blue; panels c and d), and PG (green; panels e and f). In Fig. 1c, dark colors indicate a smaller *R* values. *D* value for 160 mg/mL BSA in bulk systems was *D*_bulk_ = 25.3 ± 3.8 μm^2^/sec (Ave. ± S. D., *n* = 30). Similar *D* values in bulk systems and droplets were obtained when the TKM was used instead of PBS (Fig. S2; Table S2). Under all lipid conditions, *D*/*D*_bulk_ decreased from unity (shown in broken lines) to below unity when *R* fell below 30 μm to a certain threshold, approximately 15–20 μm. We previously reported similar *R*-dependent slow diffusion in cell-size lipid droplets containing high concentrations of linear polymer polyethylene glycol (11) and branched polymer dextran (14, 15). These experimental results clearly show that the synergistic effects of (ii) cell-size membrane confinement and (i) macromolecular crowding markedly slow down molecular diffusion in droplets than in bulk systems.

**Figure 2.**
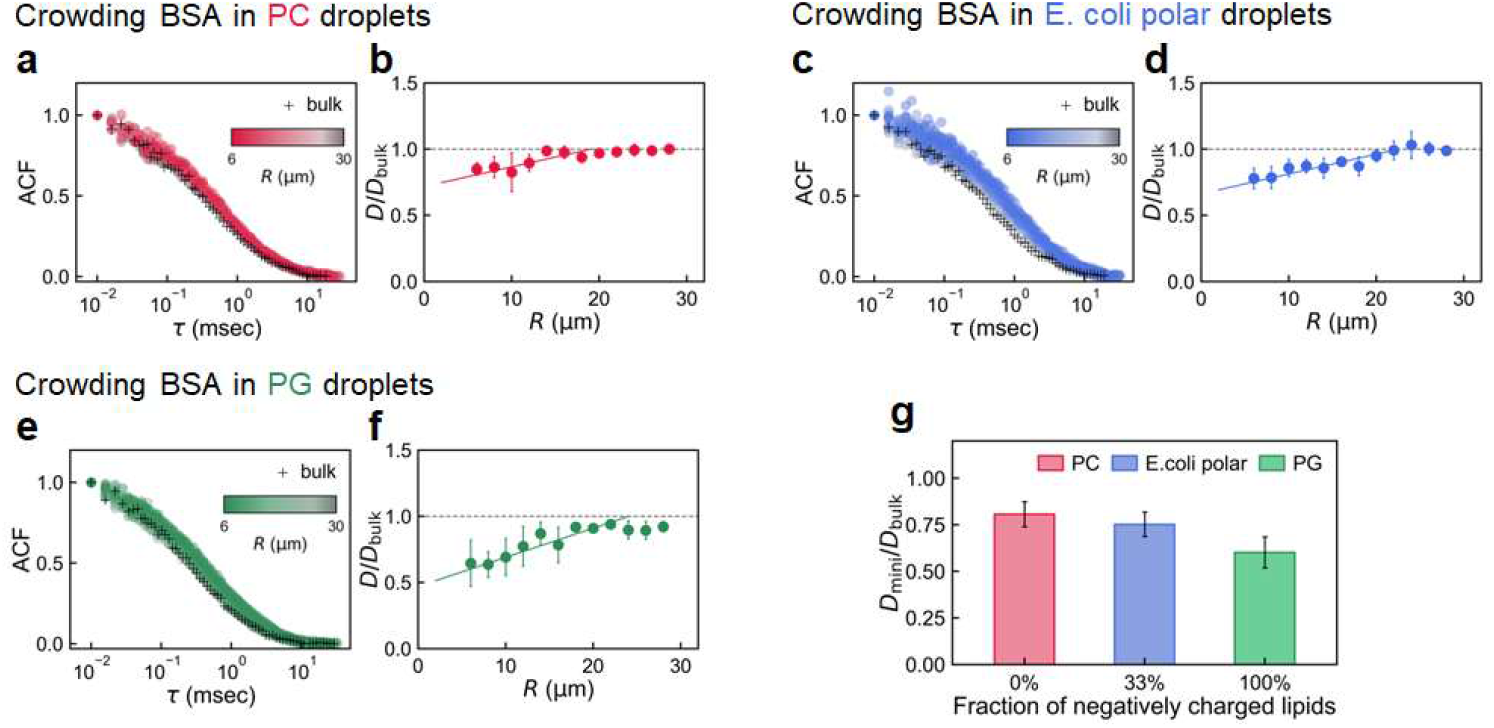
(a–f) Diffusion of TR-BSA in crowding BSA solutions within lipid droplets with *R* = 6–30 μm (different colors) and in bulk systems (black). The lipids covering the droplets are (a and b) PC, (c and d) E. coli polar, and (e and f) 1,2-dioleoyl-sn-glycero-3-phospho-(1’-rac-glycerol) (PG). Number of droplets in each graph is *n* ≥ 40, and error bars represent the S. D. Left and right panels indicate the ACF, and diffusion coefficients normalized by the corresponding bulk value, *D*/*D*_bulk_, respectively. Broken and solid lines in the right panels (b, d, and f) indicate the unity and linear fitting for small *R*, respectively. (g) Lipid-dependence of *D*_mini_/*D*_bulk_, where *D*_mini_ is the *D* value at *R* = 6 μm estimated via linear fitting. Error bars represent the standard error (S. E.) considering the fitting error.

To qualitatively evaluate slow diffusion and lipid-dependence, we fitted the *R*-dependent changes in *D* using a linear relationship (see Section S1 in SI for the analytical method). From the linear fitting (solid lines in Figs. 2b, d, and f), we estimated diffusion coefficient at minimum *R* = 6 μm, *D*_mini_ (Table S3). The normalized values of *D*_mini_/*D*_bulk_ were in the following order: PC (electrically neutral) > E. coli polar > PG (Fig. 1g). The order of membrane potential for these lipids is PC > E. coli polar > PG, which is consistent with the experimental zeta potential data (Section S2). These results suggest that the negatively charged membrane covering the droplets enhances the degree of *R*-dependent slow diffusion compared to that in bulk systems. Lipid-dependency was not clearly observed when the TKM was used instead of PBS (Fig. S3 and Table S4 for the relationship between *D*_mini_/*D*_bulk_ value and zeta potential of the lipid membrane), possibly because Mg^2+^ in the TKM weakened the effect of the negatively charged membrane.

### 2-2. Small-angle X-ray scattering (SAXS) analysis of the BSA solution in bulk systems

Before examining the synergistic effects of (i) macromolecular crowding and (ii) cell-size membrane confinement on molecular diffusion, we investigated whether the crowded environment of BSA induces clustering in bulk systems using Small-angle X-ray scattering (SAXS). This was based on the possibility that (i) macromolecular crowding increased the apparent size via clustering and hindered diffusion. To verify this, we analyzed the structure of BSA at 10 and 160 mg/mL in PBS and TKM. Figure 3 shows the SAXS intensity functions *I*(*Q*) for BSA at 10 and 160 mg/mL in (a) PBS and (b) TKM. For analysis, data at 10 mg/mL were fitted using the ellipsoidal form factor *P*(*Q*), whereas structure factor *S*(*Q*) of the screened Coulomb potential model *S*(*Q*)*P*(*Q*) was applied for the date at 160 mg/mL. From the fitting results, we determined the shape (short and long axes) and effective radius of BSA.

**Figure 3.**
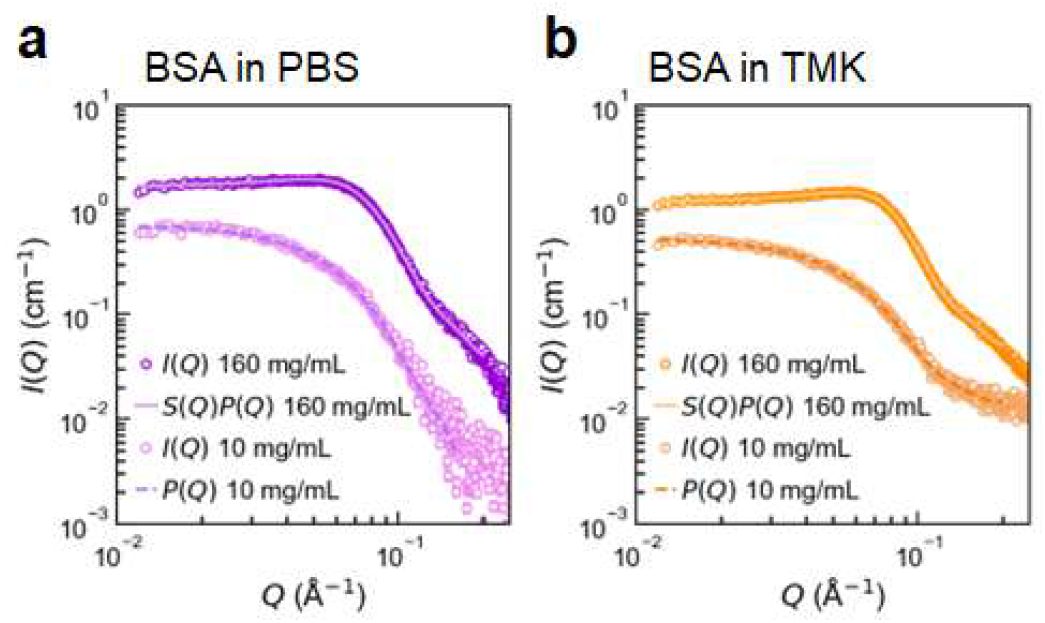
SAXS intensity functions *I*(*Q*) for 10 and 160 mg/mL BSA conditions in (a) PBS and (b) TKM. Fitting functions under 10 and 160 mg/mL BSA were the ellipsoidal form factor *P*(Q) and *P*(Q) plus screened Coulomb potential model *S*(*Q*)*P*(*Q*), respectively.

Short and long axes of the BSA molecule obtained from the ellipsoidal fitting were 20 and 44 Å, respectively, at 10 mg/mL and 20 and 40 Å, respectively, at 160 mg/mL, irrespective of the buffer type. The corresponding effective radii *R*_eff_ at 10 mg/mL (35–36 Å) were slightly higher than those at 160 mg/mL (32–33 Å). These values are consistent with the previously reported short and long axes of BSA (17 and 42 Å, respectively) with *R*_eff_ = 33 Å at 10 mg/mL BSA in 300 mM NaCl (10). The estimated volume fractions of BSA were approximately 1 and 7% at 10 and 160 mg/mL BSA, respectively. The charge parameters determined by *S(Q)* fitting were 26.4 and 18.8 at 160 mg/mL in PBS (containing 137 mM NaCl) and TKM (containing 150 mM KCl), respectively. Importantly, BSA shape remained nearly unchanged under different buffer conditions and BSA concentrations, without any cluster formation (Fig. S4).

### 2-3. Slow diffusion mechanism in cell-size spaces and its verification using ATP

The Stokes–Einstein law states that the molecular diffusion *D* of a spherical particle is inversely proportional to the solution viscosity *η* (*D* = *k*_B_*T*/6π*ηa*, where *T* is the absolute temperature, and *a* is the particle radius). The viscosity *η* of a protein solution is minimal at a the isoelectric point (pI) specifically near pH 4.7 for BSA and increases as the solution becomes more acidic or basic relative to this pI due to the electrorheological effects (16, 17). Therefore, low viscosity is possibly observed when the repulsive potential *U*_repl_ acting on the negatively charged BSA is weak. As shown in Figure 4a, *U*_repl_ involves short-range steric repulsion (normalized molecular distance, *r* < 1) and long-range electrostatic repulsion (*r* > 1), the strength of these repulsions is expressed as |*K*_2_| (see Section S3 for the details). Salt addition reduces the repulsive forces (Fig. 4b, top). Adding salt to the solution decreases *U*_repl_(*r*), which acts on the negatively charged BSA. A previous study reported that the BSA solution viscosity decreases with the addition of salt (18).

**Figure 4.**
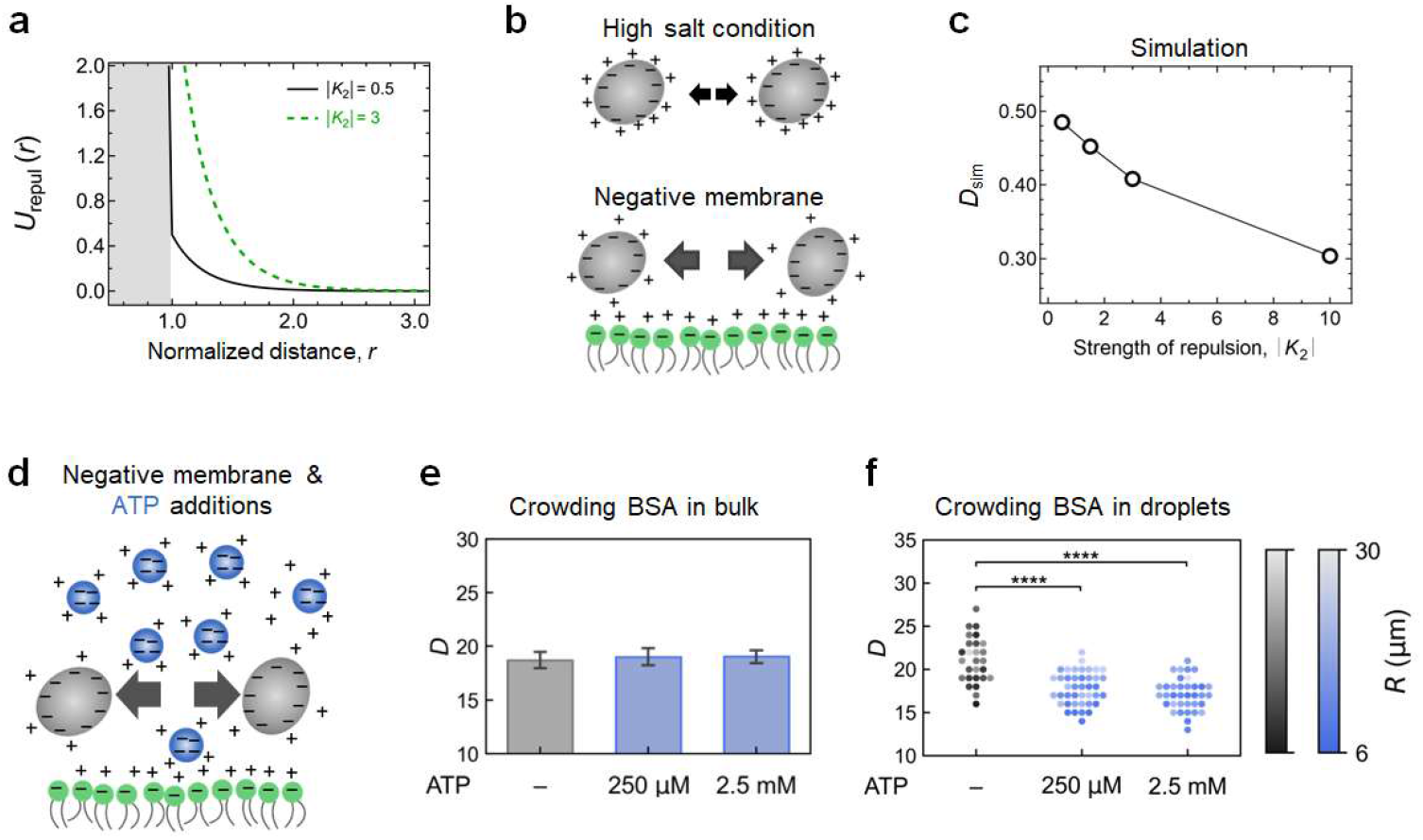
(a) Repulsive potential acting on BSA surface as a function of the normalized distance *r* from the BSA surface, considering the short-range steric and long-range electrostatic repulsions. Solid and dotted lines indicate the cases where |*K*_2_| = 0.5 and 3 for weak and strong electrostatic repulsive forces, respectively. Gray area (*r* < 1) represents the region of short-range steric repulsion. (b) Schematic diagram showing the mechanisms by which the coexisting cations and negatively charged membrane (green) alter the electrostatic repulsions in BSA. (c) Relationship between particle diffusion coefficient *D*_sim_ (dimensionless parameter) and interparticle electrostatic repulsion |*K*_2_| estimated via two-dimensional particle simulation (also see section S3 in SI). (d) Schematic diagram showing the mechanisms by which tetravalent anion ATP (shown in blue particles) and negatively charged membrane enhance the electrostatic repulsions in BSA. (e and f) Diffusion coefficient of TR-BSA *D* in crowded BSA solutions in bulk systems (a) and droplets (6 ≤ *R* ≤ 30 μm) (b) with or without ATP. Asterisks indicate the significant differences determined via Welch’s test (**** *p* < 0.0001). Color in (f) indicates the droplet radius *R* (also see Fig. S8 for *R*-*D* plots).

To investigate the mechanisms by which repulsive forces impact molecular diffusion, we performed a two-dimensional particle simulation by changing repulsion |*K*_2_| values against a fixed attractive force (Section S3). The potentials acting on the particles included short-range van der Waals attraction, short-range steric repulsion, and long-range electrostatic repulsion due to the electric double layer of the counterions (Fig. S5). As |*K*_2_| increased, estimated *D*_sim_ from the simulation decreased (Figs. 4c, S6 and S7), indicating that strong electrostatic repulsion hinders diffusion.

These simulation results can be explained by the competition for cations between BSA and the membrane. The membrane strongly attracted cations, reducing the availability of cations shielding from the negative charge of the BSA molecules (Fig. 4b, lower part). Consequently, *U*_repl_(*r*) increased. Additionally, number ratio of cations attracted to the membrane and the BSA increased as the droplet size decreased (as the membrane area-to-volume ratio increased). This possibly contributed to the considerable increase in *U*_repl_(*r*) and decrease in *D* values in the cell-size spaces.

The above-mentioned mechanisms of diffusion suppression suggest that the addition of tetravalent anion ATP further decreases BSA diffusion more. This is because ATP reduces the number of available cations to shield from the negative charge of the BSA molecules (Fig. 4d). First, we examined diffusion for crowding BSA conditions in bulk systems (160 mg/mL BSA in TKM; see section 4-2 in Materials and Methods). Addition of 250 μM or 2.5 mM ATP did not alter diffusion in bulk systems compared to that in the systems without ATP (Fig. 4e). However, ATP addition to the cell-size system (droplet with *R* ≤ 30 μm) drastically hindered diffusion. The *D* value consistently decreased as the ATP concentration increased from 250 μM to 2.5 mM, while the dependence on *R* became less pronounced (Fig. S8). Therefore, although cation accumulation on the negatively charged molecules can be disregarded in large bulk systems, it cannot be ignored in cell-size systems with membranes as it enhances the electrostatic repulsions between proteins.

### 2-4. Min protein waves facilitate BSA diffusion in cell-size spaces

Various active molecular systems affect molecular diffusion in living cells (19, 20). This study used a reaction–diffusion system for the Min protein (21, 22) to investigate the impacts of active molecular systems on the observed slow diffusion of BSA within a cell-size space. The Min wave indicates the movement of molecular populations along the cell membrane surface when the reaction between MinD and MinE is balanced. The minimum wave-generation mechanism is shown in Figure 5a. Briefly, MinD binds to the membrane in an ATP-dependent manner and gets recruited to the membrane in a positive feedback loop. MinE, which senses membrane-binding MinD, also binds to the membrane and stimulates the adenosine triphosphatase (ATPase) activity. Hydrolysis of ATP bound to MinD triggers the dissociation of MinD from the membrane. Figure 5b shows the fluorescence images of the spatial distribution of MinD (green) and MinE (not visible) in the droplets covered with the E. coli polar lipids, which also contain 2.5 mM ATP and 160 mg/mL BSA (red) in the TKM (see section 4-3 in Materials and Methods for the experimental details). Without MinE, all MinD proteins are localized to the membrane (Fig. 5b, left). In the presence of excess MinE, MinD dissociated from the membrane and was homogeneously distributed within the droplets (Fig. 5b, right). Balanced membrane binding and dissociation of MinD (MinD:MinE = 1:1 molar ratio) generated the Min protein waves on the membrane (Fig. 5b, center; Fig. 5c).

**Figure 5.**
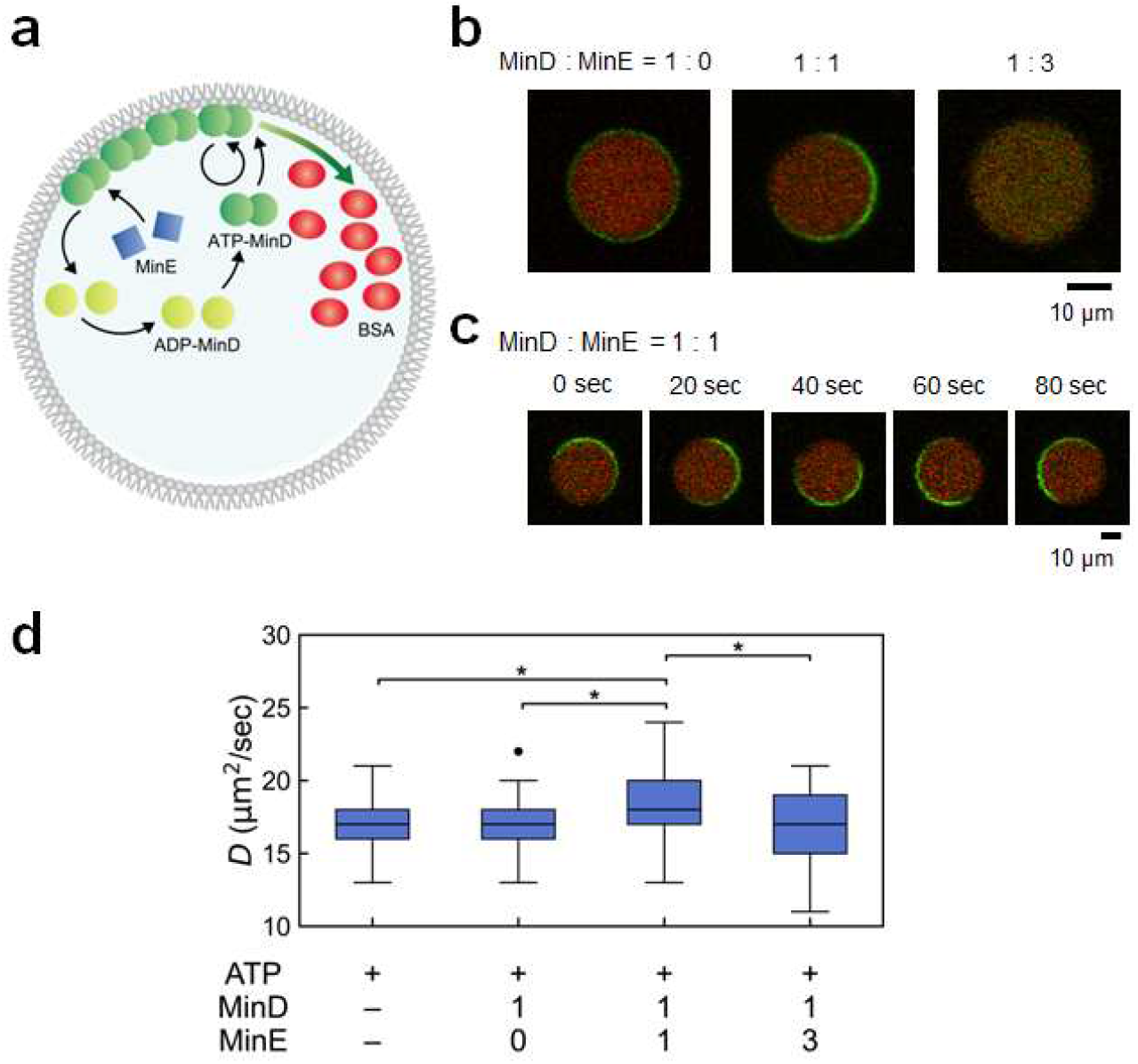
(a) Schematic illustration of the Min system in droplets. (b and c) Fluoresce images showing the (b) localization and (c) traveling wave of MinD proteins (green) on membranes with or without MinE proteins in the presence of ATP and BSA (red). Min wave is only observed at MinD:MinE = 1:1 [μM]. (d) Diffusion coefficient *D* values of TR-BSA in droplets under crowded BSA conditions with 2.5 mM ATP and various concentrations of MinE and MinD proteins. Asterisks indicate the significant differences determined via Welch’s test (* *p* < 0.05), with false discovery rate (FDR) < 0.1 for multiple comparisons. Variances at MinD:MinE = 1:1 and 1:3 [μM] were more significant than those at 0:0 and 1:0 [μM] (Table S6 and S7).

BSA diffusion was also examined at the center of the droplets (*R* ≤ 30 μm) under four different Min protein conditions: MinD:MinE = 0:0, 1:0, 1:1, and 1:3. In the absence of Min protein waves (MinD:MinE = 1:0 and 1:3), diffusion coefficient *D* remained unchanged compared to that under the conditions without any Min proteins (MinD:MinE = 0:0; Fig. 5d, left). In contrast, under Min protein wave propagation (MinD:MinE = 1:1), *D* exhibited a slight but significant increase (*p* < 0.05), which was not observed under the other three conditions. This difference was more pronounced when comparing with the effective diffusion coefficient *D*_α_ for anomalous diffusion (Section S4 and Fig. S9).

There are four mechanisms to explain the observed acceleration of BSA diffusion by Min protein waves: (1) Electrostatic screening of the membrane due to MinD membrane binding, (2) ATP depletion due to the hydrolytic activity of MinD, (3) binding and dissociation of MinD with MinE on the membrane, and (4) propagation of MinD protein waves on the membrane. Similar *D* values for MinD:MinE = 1:0 and 1:3 ruled out mechanism (1) because MinD proteins accumulated on the membrane only at MinD:MinE = 1:0. Mechanism (2) was also excluded as enzymatic ATP hydrolysis by MinD was very slow (approximately 0.1 μM/sec), and ATP concentration (2.5 mM) remained nearly constant during the measurements. Furthermore, the slight change in *D* at MinD:MinE = 1:3, which facilitated the binding and dissociation of MinD from the membranes without Min wave generation, ruled out mechanism (3). Therefore, mechanism (4), i.e., propagation of Min proteins along the membrane, is the most plausible explanation for the accelerated diffusion exclusively observed at MinD:MinE = 1:1. Notably, variances in *D* were considerably larger at MinD:MinE ratios of 1:1 and 1:3 compared to those at other ratios (MinD:MinE = 0:0 and 1:0; Fig. 5d; Table S6 and S7). This increased variability suggests that mechanism (3), *i*.*e*., balance between MinD membrane binding and dissociation in the presence of MinE, affects the fluctuation in *D* values rather than its average value.

## 3. Discussion

### 3-1. Protein diffusion regulation by cell-size membrane confinement

Under macromolecular crowding conditions, BSA diffusion was regulated by membrane confinement in cell-size spaces (Fig. 2). Simulations showed that the increased electrostatic repulsion between BSA molecules inhibited diffusion (Fig. 4c). Under physiological conditions, both BSA and the membrane were negatively charged (Fig. 2g), and membrane confinement reduced the number of available cations, thereby increasing electrostatic repulsion and inhibiting diffusion. As the spatial size decreased and membrane area-to-volume ratio increased, the membrane strongly influenced the repulsive interactions between BSA molecules.

Using green fluorescent protein (GFP) instead of TR-BSA, we previously reported that membrane confinement in cell-size spaces inhibits GFP diffusion under macromolecular crowding conditions, regardless of shape (spherical or disk shape) (11). Upper volume limit for slow diffusion is larger in the disk space than in the spherical space. Furthermore, PEGylation of membrane (15) and neutralization of the membrane surface charge (Fig. 2g) restore diffusion inhibition. These results suggest that the membrane effect of modulating repulsive interactions that determines diffusion depends on the changes in the membrane area-to-volume ratio rather than the volume or local curvature of the space.

In bulk systems, enhanced electrostatic repulsion by ATP did not alter molecular diffusion in this study (Fig. 4e). In contrast, ATP significantly inhibited diffusion in the cell-size spaces with membranes (Fig. 4f). These findings call into question the conventional assumption that slight changes in salt concentration, protein concentration, and pH, which do not affect the molecular behaviors in bulk systems, can be ignored in cell-size spaces. Similar to the multimolecular competition among various proteins for membranes in cell-size spaces (23), repulsive interactions within cells must be evaluated considering the multi-ion competition among various ions for membranes.

### 3-2. Regulation of repulsive interactions in living cells

In colloidal systems, strong attractive or repulsive interactions lead to a glassy state characterized by extremely slow diffusion. This glassy state is observed in the cytoplasm of metabolically inactive bacteria (24, 25). Decreased ATP levels and a drop in cytoplasmic pH < 7 often trigger such inactivity (26), and these changes hinder the diffusion of proteins with a pI of approximately 4 (4). Our findings suggest that electrostatic repulsive interactions are crucial for protein diffusion regulation in living cells (Figs. 2 and 4). Anions, such as ATP, help to regulate these repulsive interactions. As ATP levels increase, protein solutions initially aggregate due to reduced electrostatic repulsion. Accordingly, the aggregated state transitions to a dispersed state with slower molecular diffusion due to excess repulsion and then to a reaggregation state due to hydrophobic interactions (27-29). This reentrant behavior suggests that cells maintain cytoplasmic fluidity by carefully regulating the concentrations of ATP (and other anions) and proteins, achieving an appropriate balance of repulsive interactions. This control of repulsive interactions should involve the regulation of cell membrane surface charge (30, 31).

Here, our results suggest that active molecular systems along membranes, such as the Min protein waves, further control such repulsive interactions (Fig. 5). A simple explanation for this control is that the Min protein wave propagation dynamically alters the surface charge and effective size of certain molecules, thereby facilitating their diffusion (32). Both the static and spatiotemporal control of repulsive interactions are important to regulate molecular behaviors. Future studies should explore the mechanisms by which dynamic molecular systems affect these repulsive interactions to modulate the molecular behaviors in living cells.

In summary, we found that the negatively charged membrane significantly hindered protein diffusion inside cell-size space due to enhanced electrostatic repulsion caused by the reduced cation availability. Notably, ATP further inhibited protein diffusion in cell-size space but did not significantly affect that in bulk solution. These results highlight the crucial role of membranes in regulating the electrostatic repulsions in cells, thereby influencing various molecular behaviors, such as diffusion. Our results also showed that Min protein wave propagation on the membrane actively counteracted the excessive repulsive interactions, indicating an active regulatory mechanism fine-tuning intracellular molecular diffusion in cells.

## 4. Materials and Methods

### 4-1. Materials

Globular protein BSA from Sigma-Aldrich (A6003; Saint Louis, MO, USA) was used to mimic the macromolecular crowding conditions in the cytoplasm. Fluorescence-labeled BSA (TR-BSA A2307; Invitrogen, Carlsbad, CA, USA) was used to measure BSA diffusion via FCS. Additionally, 5-carboxy tetramethylrhodamine (TAMRA) with a diameter of 1.7 nm was used for calibration. 1,2-dioleoyl-sn-glycero-3-phosphocholine (PC; 850375C), 1,2-dioleoyl-sn-glycero-3-phospho-(1’-rac-glycerol) (sodium salt; PG, 840475C), and *E. coli* extract polar (E. coli polar; 100600C) were purchased from Avanti Polar Lipids Inc. (Alabaster, AL, USA). Lipid composition of E. coli polar included approximately 67% phosphatidylethanolamine, 23% phosphatidylglycerol, and 10% cardiolipin. Except for BSA, these materials were used without further purification. As lipid solvents, chloroform (Fujifilm Wako Pure Chemical, Osaka, Japan) and mineral oil (kinetic viscosity of 14–17 mm^2^/sec at 38 °C; Nacalai Tesque, Kyoto, Japan) were treated with molecular sieves. Two types of buffers were used to prepare the BSA solutions: PBS (pH 7.4; 314-90185; Nippon Gene Co., Ltd., Toyama, Japan), containing 137 mM NaCl, 2.68 mM KCl, 8.1 mM Na_2_HPO_4_, and 1.47 mM KH_2_PO_4_, and TKM prepared by mixing 25 mM Tris-HCl (pH 7.6; Nacalai Tesque), 150 mM KCl, and 5 mM MgCl_2_ (Fujifilm Wako Pure Chemical).

### 4-2. Preparation of BSA solution

BSA molecules were dissolved in PBS, and BSA solution was washed off before use. Briefly, 100 mg of BSA was dissolved in 1 mL of PBS. It was washed with a 10-fold volume of PBS using 30 kDa filter units (Amicon Ultra-0.5; Merck Millipore, Darmstadt, Germany) and a centrifuge (Universal Refrigerated Centrifuge Model 6200; Kubota Corp., Japan) at 4 °C. After washing 4–5 times, BSA sample was concentrated to 250–450 mg/mL and stored at 25 °C until use. For FCS measurement, BSA in PBS was mixed with 120 nM TR-BSA. BSA concentration was measured using BioSpec-nano (Shimadzu Corp., Kyoto, Japan). The BSA solution was further used for the Min protein systems by replacing PBS with TKM. Nominal diameter of BSA is 7–8 nm (33). Under 10 and 160 mg/mL BSA conditions, the estimated volume fractions were approximately 1 and 11%, respectively (see SI of our previous report (11) for analysis). Moreover, intermolecular distance between BSA for 160 mg/mL BSA was approximately 2 nm, which was much smaller than the diameters of BSA and TR-BSA (TR size = approximately 1.5 nm (34)).

### 4-3. Preparation of cell-size droplets

Based on our previous report on the slow diffusion within BSA droplets (11), we used a 160 mg/mL BSA solution in this study. The samples were prepared in PBS or TKM solutions containing 120 nM TR-BSA. Then, 1 mg/mL lipid of PC in mineral oil was prepared by mixing lipid-in-chloroform with mineral oil and evaporating the chloroform at 70 °C overnight. For E. coli polar or PG, lipid-in-chloroform was dried into a thin film inside a glass tube. Subsequently, mineral oil was added, and the mixture is stirred, followed by ultrasonication at 60 °C and 40 kHz for 90 min using the ultrasonic cleaner (MCD-6P; As One, Osaka, Japan.) The BSA solution was added to the lipid-oil solution at a volume ratio of 1:50 (w/o). By emulsification via pipetting, we prepared droplets with a radius (*R*) of 6−30 μm. Subsequently, the droplet solution was sandwiched between two cover glasses (No. 1; 0.12– 0.17 mm; Matsunami Glass Co., Osaka, Japan) with spacers of approximately 0.1 mm thickness (double-sided sticky tape; J0410; Nitoms, Inc., Tokyo, Japan).

### 4-4. Preparation of Min system

Min proteins (His-msfGFP-MinC, His-MinD, and MinYE-His) were expressed and purified as previously described (21, 22). The chemical reagents used in this section are the same as those used in previous publication (35). Briefly, *E. coli* BL21-CodonPlus(DE3)-RIPL cells were transformed with pET15-msfGFP-MinC (22), pET15-MinD (21), or pET29-MinE (21) and cultivated in the Luira– Bertani medium with 0.1 mg/mL ampicillin (for msfGFP-MinC and MinD) or kanamycin (for MinE) at 0.025 mg/mL. After optical density at 600 nm reached 0.8, Min proteins were expressed by the addition of 1 mM isopropyl-β-D-thiogalactopyranoside. The cells were further cultivated at 37 °C for 3 hours and harvested. Then, the cells were resuspended in a lysis buffer (50 mM Tris-HCl (pH 7.6), 300 mM NaCl, 1 mM phenylmethylsulfonyl fluoride (PMSF), 20 mM imidazole, and 1 mM dithiothreitol) and sonicated using Sonifier250 (Branson, Danbury, CT, USA) or Sonifier SFX150 (Branson). For MinD, 0.1 mM ADP was added to the lysis buffer, and the supernatant was collected after centrifugation of the lysate at 20,000 × *g* for 30 min at 4 °C. The supernatant was filtered using HPF Millex HV (Merck Millipore) and mixed with Ni Sepharose 6 Fast Flow (Cytiva, Tokyo, Japan). The samples were shaken at 4 °C for 30 min and loaded onto the Poly-Prep Chromatography Column (Bio-Rad, Hercules, CA, USA). Ni-NTA beads in the Poly-Prep Chromatography Column were washed with the Ni-wash buffer (50 mM Tris-HCl (pH 7.6), 300 mM NaCl, 1 mM PMSF, 0.1 mM ethylenediaminetetraacetic acid (EDTA), 25 mM imidazole, and 10% glycerol), and His-tagged protein was eluted using the Ni-elution buffer (50 mM Tris-HCl (pH 7.6), 300 mM NaCl, 1 mM PMSF, 0.1 mM EDTA, 500 mM imidazole, and 10% glycerol). For His-msfGFP-MinC, the eluted solution was diluted six times by adding 50 mM HEPES-KOH (pH 7.6) and purified using Q Sepharose High Performance (Cytiva). Q Sepharose beads bound to His-msfGFP-MinC were washed with the Q-wash buffer (50 mM Hepes-KOH (pH 7.6) and 50 mM KCl), and msfGFP-MinC was eluted using the Q-elution buffer (50 mM Hepes-KOH (pH 7.6) and 50 mM KCl).

For all proteins, the buffer solution was exchanged to a storage buffer solution (50 mM Hepes-KOH (pH 7.6), 150 mM GluK, 0.1 mM EDTA, 10% glycerol, and 0.1 mM ADP in the case of MinD) via repeated ultrafiltration using Amicon Ultra-0.5 3K (Merck Millipore) for MinE or Amicon Ultra-0.5 10K (Merck) for msfGFP-MinC and MinD. Purified protein concentrations were estimated by quantifying the band intensities after Coomassie brilliant blue staining of sodium dodecyl sulfate-polyacrylamide gels using the Fiji software (National Institutes of Health, Bethesda, MD, USA).

### 4-5. Fluorescence observation of BSA droplets

BSA droplets with or without Min system were observed using a confocal laser scanning microscope (IX83; FV1200; Olympus Inc., Tokyo, Japan) equipped with a water-immersion objective lens (UPLSAPO 60XW; Olympus Inc.). TR-BSA and msfGFP-MinC were excited at wavelengths of 559 and 473 nm and detected in the ranges of 575–675 and 490–540 nm, respectively. The fluorescence images were subsequently analyzed using Fiji software.

### 4-6. Translational diffusion analysis

Translational diffusion of TR-BSA was analyzed using point FCS (FV1200 confocal microscope integrated with a molecular diffusion package; Olympus Inc.), as described in our previous studies (11, 14). A water-immersion objective lens (UPLSAPO 60XW; Olympus Inc.) was used to perform point FCS with minimal laser power (approximately 0.3 μW) to minimize the temperature change caused by laser irradiation. To eliminate the effect due to the high refractive index of oil phase and concentrated BSA solution, diffusion was performed within droplets with *R* > 5 μm at a z-height of 5 μm. Droplets partially adhered to the cover glass, as shown in Fig. 1a, were selected. The height of the adhered droplets with *R* = 10 μm is higher than 5 μm. Autocorrelation functions (ACFs) were derived using the microscope-accompanying software. Illumination volume was calibrated using the TAMRA solution (20 nM) with the same diffusion coefficient value of 280 μm^2^/s. For TR-BSA, *x–y* radius of the illumination volume *w*_0_ and its ratio *s* to its *z*-height *w*_z_ (*s* = *w*_z_/*w*_0_, structure factor) were 0.19–0.22 μm and 4–6, respectively. Translational diffusion behaviors were analyzed by fitting ACFs as a function of correlation time *τ*:

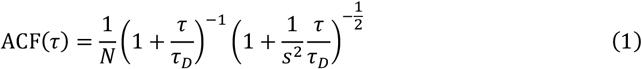

where *N* is the number of fluorescent molecules, and *τ*_D_ is the characteristic decay time. Using *τ*_D_ and *w*_0_, diffusion coefficient *D* was derived as follows:

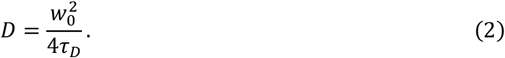

### 4-7. Small-angle X-ray scattering (SAXS) experiment

To compare the structures of 10 and 160 mg/mL BSA in PBS and TKM, we analyzed them by using Small Angle X-ray Scattering (SAXS). The solutions were prepared as mentioned above section 4-2 and were filtered through a 0.2 μm pore size filter (6784-2502; Whatman; Cytiva) before measurement. A laboratory SAXS instrument (SAXSpoint 2.0; Anton Paar GmbH, Graz, Austria) was used. Incident X-ray wavelength was 1.542 Å of Cu Kα radiation. The scattering patterns were detected using a 2-dimensional hybrid pixel detector with a spatial resolution of 75 μm (EIGER R 1M, DECTRIS Ltd., Barden, Switzerland). The sample-to-detector distance was 560 mm. Wavenumber *Q* = 4πsin(*θ*/2)/*λ*, where *θ* is the scattering angle, and *λ* is the X-ray wavelength. The data were corrected in the *Q* range of 0.01–0.3 Å^-1^. All the measurements were performed in a vacuum chamber. The samples were sealed in a measurement cell with 30 μm thick cover-glass windows. The thickness of each sample was 1 mm. The cells were mounted on a temperature-controlled stage in the vacuum chamber.

SAXS intensity functions *I*(*Q*) for 10 or 160 mg/mL BSA solutions were fitted with the ellipsoidal form factor *P*(Q) and structure factor *S*(Q) of the screened Coulomb potential model *S*(*Q*)*P*(*Q*) (10, 36) using the SasView software (37). First, we analyzed the ellipsoidal shape with *P(Q)*. Short and long axes of the BSA molecule (*r*_pol_ and *r*_equ_, *r*_pol_ < *r*_equ_) were 20 and 44 Å. respectively for 10 mg/mL, and 20 and 40 Å, respectively, for 160 mg/mL, regardless of the buffer. Effective radius *R*_eff_ was estimated considering the ellipsoidal shape (38).

## ASSOCIATED CONTENT

### Supporting Information

Supporting Information is freely available for this article. Fitting of R-dependent changes in diffusion coefficient *D* (section S1), linear relationship between the normalized diffusion coefficient and membrane charge determined by the zeta potential (section S2), numerical simulation of circular particles diffusing on a two-dimensional plane (section S3), translational analysis of anomalous diffusion (section S4) are described in the Supporting Information Text. Autocorrelation function (ACF) and *R*-dependence of *D* in 1,2-dioleoyl-sn-glycero-3-phosphocholine (PC) droplets in the non-crowding (Fig. S1) and crowding (Fig. S2) bovine serum albumin (BSA) solutions in the TKM for three types of lipids, effects of buffer type and membrane charge (Fig. S3), structure factor determined via small-angle X-ray scattering (SAXS) intensity analysis (Fig. S4)and potential energy of BSA particles (Fig. S5) on numerical simulation, relationships between mean square displacement (MSD) and repulsive interactions (Fig. S6) and dependence of *D*_sim_ and repulsion strength (Fig. S7) during numerical simulation, *R*-dependence of diffusion coefficient *D* on ATP concentration (Fig. S8), and diffusion coefficient considering anomalous diffusion of BSA with the Min system (Fig. S9) constitute the Supporting Information Figures. Diffusion coefficients of bulk and PC droplets under non-crowding BSA conditions (Table S1) and bulk values in PBS or TKM (Table S2), fitting results of the linear relationships between each buffer and lipids (Table S3), zeta potentials of BSA and lipid membranes (Table S4), zeta potentials of BSA with ATP (Table S5), diffusion coefficients under crowding conditions in the presence of the Min system (Table S6), and F-test results of the variances of diffusion coefficients (Table S7) constitute the Supporting Information Tables.

### Notes

The authors declare no competing financial interest.

## Acknowledgements

This research was partially funded by the Japan Society for the Promotion of Science (JSPS) KAKENHI (grant numbers 25K17355 (H.S.), 22H01188, 24H02287 (M.Y.), 22K19299, 22H04851 (K.F.)), the Ishii-Ishibashi Fund (The Keio University Grant for Early Career Researchers)(K.F.), and the Japan Science and Technology Agency (JST) (grant numbers FOREST, JPMJFR213Y; CREST (JPMJCR22E1) (M. Y.)). The SAXS experiment was carried out by the JRR-3 joint user program managed by the Institute for Solid State Physics, the University of Tokyo (Proposal No. 24404).

