## Supplemental information for "Membrane-enhanced repulsive interactions regulate protein diffusion in cell-size space"

### S1. Linear relationship between diffusion and radius in small droplets

To quantify the $R$-dependent change of the diffusion coefficient $D$ inside droplets (Fig. 2, a-f), we fitted the relationship using the following equations:

$$D = \begin{cases} l(R - R^*) + D^* & R \leq R^* \\ D^* & R > R^* \end{cases}. \quad \text{(S1)}$$

When the droplet radius $R$ is larger than a specific size $R^*$, $D$ is independent of $R$. When $R$ is smaller than $R^*$, $D$ decreases as $R$ becomes smaller, and the degree of this slow diffusion is expressed with the parameter $l$. For the fitting, the value of $D^*/D_{\text{bulk}}$ was fixed to be unity.

### S2. Zeta potential analysis of BSA and lipid membranes and its relationship to BSA diffusion

The zeta potential of lipid membranes was measured using small unilamellar vesicles (SUV) about 100 nm in diameter, prepared from a chloroform solution containing 10 mg/mL of lipid (PC, E. coli polar lipid, or PG). Nitrogen gas was used to evaporate the chloroform, yielding dry lipid films on a glass tube. PBS and TKM were then added for hydration, resulting in a 1 mg/mL vesicle solution, which was ultrasonicated for 30 minutes and extruded through a 100 nm polycarbonate film 21 times to prepare monodisperse SUVs.

Zeta potentials of SUVs were measured using a zeta-potential meter (Zetasizer Nano ZS Zen 3600, Malvern Instruments Ltd., Worcestershire, UK) with a disposable cell (DTS1070, Malvern Panalytical Ltd., Worcestershire, UK). Each SUV solution was diluted to 0.1 mg/mL before the measurement. For BSA, zeta potentials were measured with a zeta-potential meter (ELSZneo, Otsuka Electronics Co., Ltd., Osaka, Japan) with a flat electrode cell unit (Otsuka Electronics Co., Ltd.).

To explore the relationship between the $R$-dependent slow diffusion and membrane surface charge, we measured the zeta-potential $\zeta$ for small liposomes (see Materials and Methods section 4-8) composed of PC, E. coli polar, and PG in PBS or TKM. Under all conditions, the membrane exhibited negative $\zeta$ values, irrespective of the lipid composition or buffer type. The magnitude of $\zeta$ value, $|\zeta|$ followed the order PG > E. coli. Polar > PC, indicating PG content increases the negative membrane charge (*1*). Additionally, the presence of $Mg^{2+}$ in TKM reduces the $|\zeta|$ compared to PBS for all lipid types (*i.e.*, $|\zeta_{\text{PBS}}| > |\zeta_{\text{TKM}}|$). This trend was also observed for BSA in bulk. The $\zeta$ value of 10 mg/mL BSA in TKM ($\zeta_{\text{PBS}} = -20.1 \pm 0.7$ mV; Ave. ± S. D., $n = 5$) was smaller than that in PBS ($\zeta_{\text{TKM}} = -17.4 \pm 1.3$ mV, $n = 3$). This reduction in $|\zeta|$ suggests that divalent $Mg^{2+}$ in TKM are more effective in screening the surface charges of BSA and membranes than monovalent ions such as $K^+$ and $Na^+$. To further investigate, we plotted the normalized diffusion coefficient, $D_{\text{mini}}/D_{\text{bulk}}$ against membrane charge, $\zeta$ in Figure S4. Under the same buffer conditions, the $D_{\text{mini}}/D_{\text{bulk}}$ exhibited a linear relationship against $\zeta$, showing that the cell-size confinement within more negatively charged membrane leads to slower diffusion rates.

### S3. Simulation of diffusion behavior of BSA molecules considering repulsive interactions

Based on the experimental results, we expect electrostatic repulsion to slow BSA diffusion (Figs. 4a, b). To verify this expectation, we investigated how this repulsion affects the particles' diffusion by performing numerical simulations of circular particles diffusing on a two-dimensional plane. In our simulations, we employ the two-Yukawa potential $U_{\mathrm{TY}}(r)$, which is defined as follows:

$$U_{\mathrm{TY}}(r) = \begin{cases} 4\left(\frac{1}{r}\right)^{12} & \text{if } r < 1 \\ -K_1 \exp\frac{[-Z_1(r-1)]}{r} - K_2 \exp\frac{[-Z_2(r-1)]}{r} & \text{if } r \geq 1 \end{cases}, \tag{S2}$$

where $K_1$ and $K_2$ are normalized by $k_{\mathrm{B}}T$ , $k_{\mathrm{B}}$ is the Boltzmann constant and $T$ is the absolute temperature, and $r$ is the interparticle distance normalized by the particle diameter $\sigma$. The term $4(1/r)^{12}$ for $r < 1$ is the hard sphere potential. Positive values of $K_1$ and $K_2$ correspond to attractive interactions, whereas negative values correspond to repulsive interactions. For BSA solutions in the presence of salts, $K_1$ takes positive values, while $K_2$ takes negative values (*2*). The interaction range, characterized by the inverse of the screening parameter $Z$, determines the spatial extent of attractive or repulsive forces. For BSA solutions, $K_1 = 0.6$ and $Z_1 = 12$, and these values are independent of salt concentration and type (*2*). $Z_2$ is defined as $\sigma / \kappa^{-1}$, where $\kappa^{-1}$ is the Debye length. In this study, we set $\sigma = 1$ and varied $K_2$ and $Z_2$ as parameters to investigate changes in the diffusion behavior of the particles. Figure S5 shows the shape of $U_{\mathrm{TY}}(r)$ when $Z_2 = 1$ fixed. The red, orange, green, and blue lines indicate $K_2 = -0.5, -1.5, -3$, and $-10$, respectively. In this study, the cutoff length was set to $r = 5\sigma$.

The motion of each particle is governed by the dimensionless Langevin equation:

$$\frac{dv_i(t)}{dt} = -\xi^* v_i(t) + F_i(t) + \sqrt{\frac{2\xi^* T^*}{\Delta t}} R_{\mathrm{G}} \tag{S3}$$

where $F_i(t) = -\partial U_{\mathrm{TY}}(r_i) / \partial r_i$ represents the potential force acting on particle $i$, $\xi^* \left(= \xi\sqrt{a^2/m\,\epsilon}\right)$, where $a$ is the unit of length, $m$ is the mass of the particle, and $\epsilon$ is the unit of energy) is the ratio of the friction coefficient to the interaction strength, and $T^*(= k_{\mathrm{B}}T/\epsilon)$ is the ratio of thermal energy to interaction strength. $R_{\mathrm{G}}$is a Gaussian random number vector with a mean of 0 and a standard deviation of 1 for each component. In this study, we set $\xi^* = 1$ and $T^* = 1$. The above Langevin equation was solved using the semi-implicit Euler method:

$$v_i(t + \Delta t) = v_i(t) - \xi^* v_i(t)\Delta t + F_i(t)\Delta t + \sqrt{2\xi^* T^* \Delta t} R_{\mathrm{G}} \tag{S4}$$

$$r_i(t + \Delta t) = r_i(t) + v_i(t + \Delta t)\Delta t. \tag{S5}$$

The time step $\Delta t$ was set to 0.001. The packing fraction of the two-dimensional particles was set to $\phi = \sum_{j=1}^{N} \pi\sigma_j^2 / 4L^2 = 0.152$, which corresponds to the two-dimensional area fraction converted from the three-dimensional volume fraction of BSA in the experiment. The number of particles was set to $N = 1936$, and the simulation box size was set to $L = 100$. Periodic boundary conditions were applied.

Figure S6 shows the relationship between the mean squared displacement (MSD) of all particles and time $t$ when $Z_2 = 1$ is fixed. The red circle, orange square, green triangle, and blue inverted triangle symbols represent $K_2 = -0.5$, $-1.5$, $-3$, and $-10$, respectively. For $t > 1$, the MSD exhibits a linear dependence on time, with a slope of 1, which is characteristic of normal diffusion. The MSD decreases as $K_2$ becomes smaller at the same $t$. Figure S7 shows the dependence of the diffusion coefficient $D_{\mathrm{sim}}$, obtained by fitting MSD = $4D_{\mathrm{sim}}t$, on the repulsion strength $|K_2|$. The red circle, orange square, green triangle, and blue inverted triangle symbols represent $Z_2$=1, 3, 5, and 8, respectively. These results indicate that as the magnitude of repulsive interactions $|K_2|$ increases, particle mobility decreases, leading to a reduction in the diffusion coefficient $D$.

### S4. Translational diffusion analysis considering anomalous diffusion

We analyzed the translational diffusion behaviors by fitting ACF($\tau$) as a function of the correlation time $\tau$, with a fractional Brownian motion (fBM) model with an anomalous exponent parameter $\alpha$.

$$\mathrm{ACF}(\tau) = \frac{1}{N}\left(1 + \left(\frac{\tau}{\tau_{\mathrm{D}}}\right)^{\alpha}\right)^{-1}\left(1 + \frac{1}{s^2}\left(\frac{\tau}{\tau_{\mathrm{D}}}\right)^{\alpha}\right)^{-\frac{1}{2}} \tag{S6}$$

where $N$ is the number of fluorescent molecules, is the anomalous exponent, $\tau_{\mathrm{D}}$ is the characteristic decay time. As for Brownian diffusion mode, $\alpha$ is fixed to 1. From $\tau_{\mathrm{D}}$ and $w_0$, the diffusion coefficient $D_\alpha$ can be derived as follows:

$$D_\alpha = \frac{w_0^2}{4\tau_{\mathrm{D}}^{\alpha}} \tag{S7}$$

Figure S9 indicates the diffusion coefficient $D_\alpha$ of BSA fitted by the Equations (S6, S7).

**Supporting Information Figures**

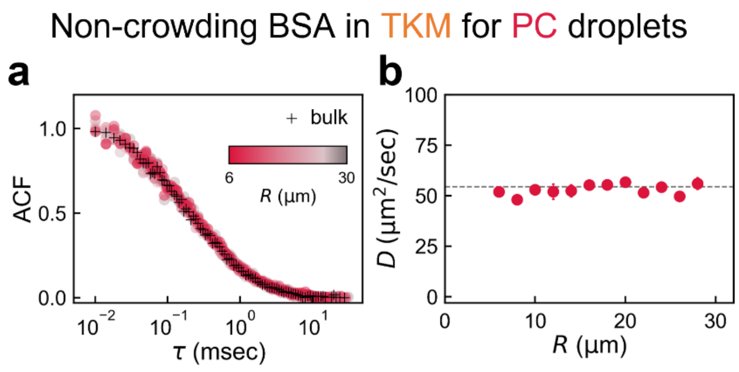


**Figure S1.** Molecular diffusion of TR-BSA inside PC droplets containing 10 mg/mL BSA in TKM. (a) ACFs for various sized droplets (red) and in bulk (black). (b) *R* dependence of diffusion coefficient *D* obtained by ACF fitting. The average value of *D* for droplets is 53.2 ± 3.4 μm$^2$/sec (Ave. ± S. D., *n* = 19). Bulk value is indicated by the dashed line ($D_{\text{bulk}}$ = 54.5 ± 1.3 μm$^2$/sec (Ave. ± S. D., *n* = 6).

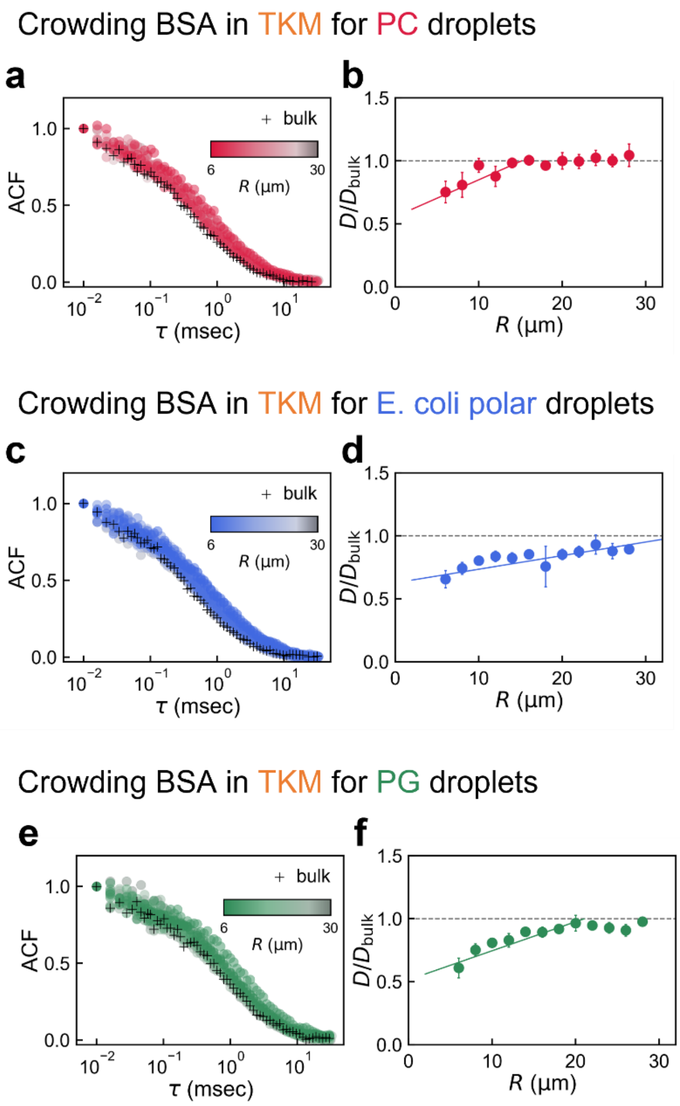


**Figure S2.** (a-f) Diffusion of TR-BSA in crowding BSA in TKM inside lipid droplets with $R$ = 6–30 µm (shown in color) and in bulk (shown in black). The lipid covering the droplets is (a, b) PC, (c, d), E. coli polar, and (e-f) PG. The number of droplets in each graph is $n \geq 40$ and the error bar is S. D. The left and right panels are ACF, and the diffusion coefficients normalized by the corresponding bulk value, $D/D_{\text{bulk}}$, respectively. The broken and solid lines in right panels (b, d, f) represent the unity and linear fitting for small $R$, respectively.

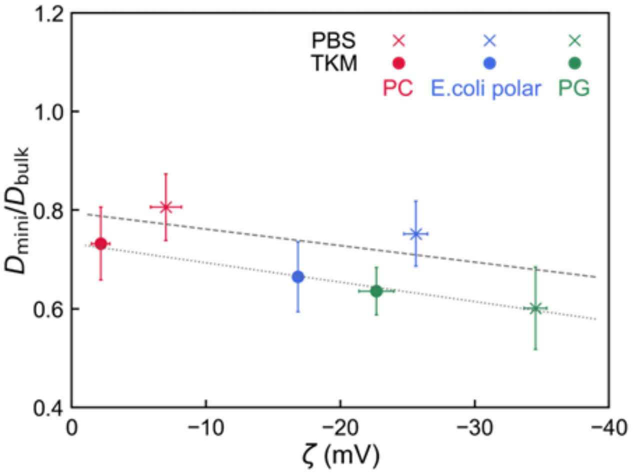


**Figure S3.** The relationship between $D_{\text{mini}}/D_{\text{bulk}}$ and the membrane surface charge $\zeta$. The $D^*$ is the estimated $D$ value at $R$ = 6 μm from the linear fitting in Figs. 2 and S2 and summarized in Table S3.

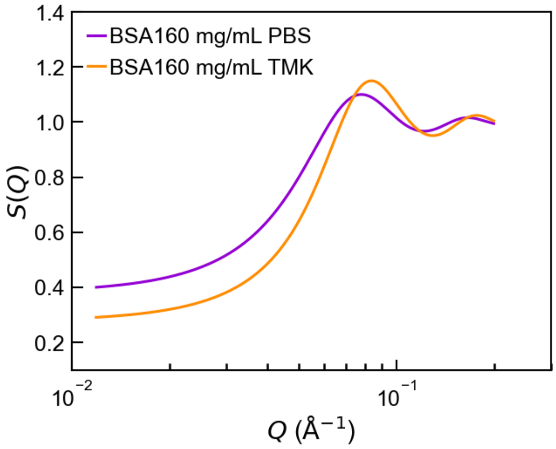


**Figure S4.** Structure factor $S(Q)$ obtained from SAXS intensity $I(Q)$ analysis results. Purple line is 160 mg/mL in PBS, orange line is TKM. Each data was obtained from the fitting of a screened Coulomb potential model $S(Q)P(Q)$ using the BSA's ellipsoidal shape with $P(Q)$ of 10 mg/mL as shown in Fig.3.

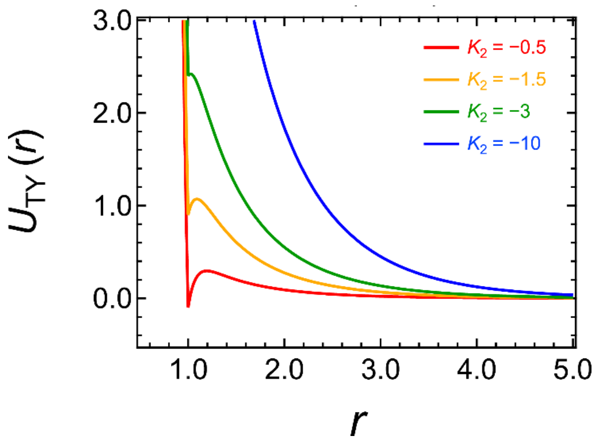


**Figure S5.** The shape of the potential energy $U_{\mathrm{TY}}(r)$ for various $K_2$ with the fixed $Z_2$ = 1. The $r$ is the interparticle distance normalized by the particle diameter, therefore $r$ = 1 corresponding to the surface of the particle.

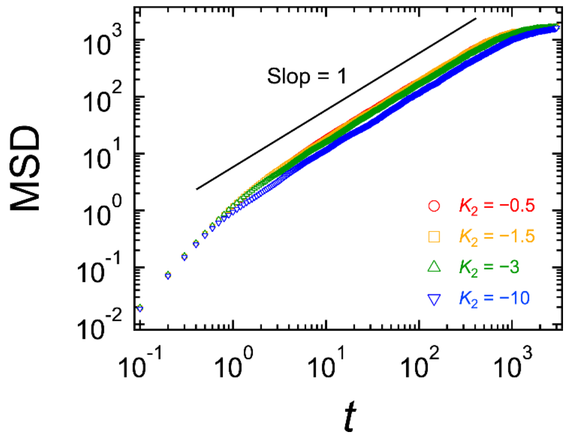


**Figure S6.** The average MSD of all particles when $K_2$ is varied with fixed $Z_2$ = 1.

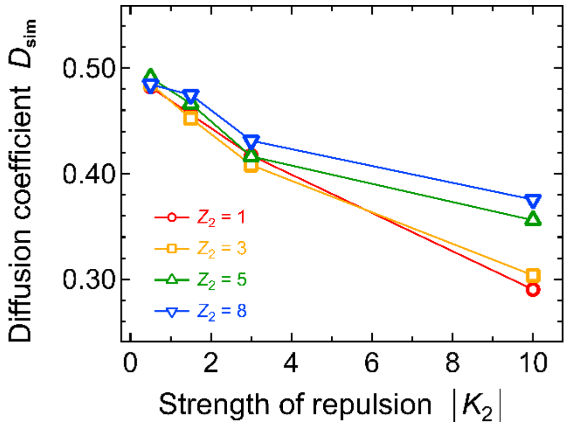


**Figure S7.** Dependence of the diffusion coefficient $D_{sim}$ on the strength of the repulsive force $|K_2|$ while maintaining a constant attractive force. The color represents the strength of the attractive force $Z_2$.

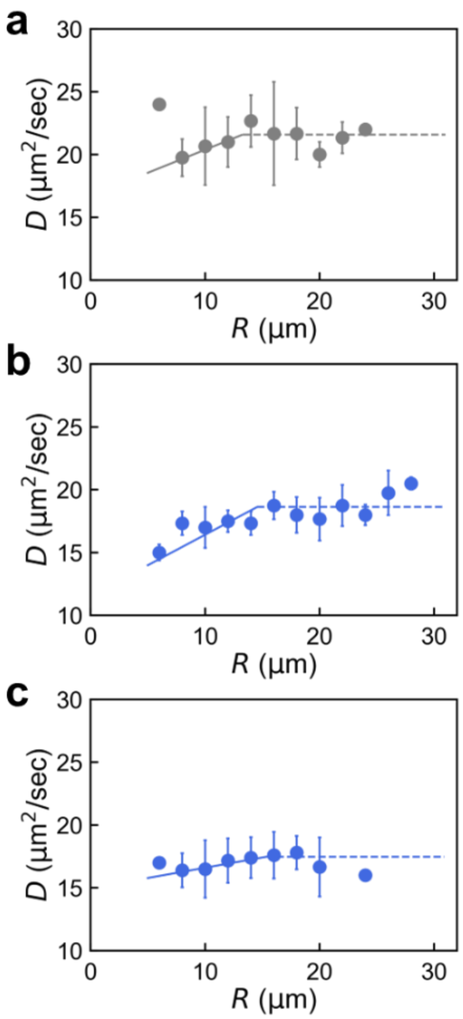


**Figure S8.** Impact of ATP addition on the R-dependent slow diffusion of TR-BSA in crowding BSA droplets. 160 mg/mL BSA in TKM is confined within E. coli polar droplets under three different conditions: without ATP (a), with 250 μM ATP (b), and with 2.5 mM ATP (c). The solid and dashed lines represent the fitting results based on Eq. (S1).

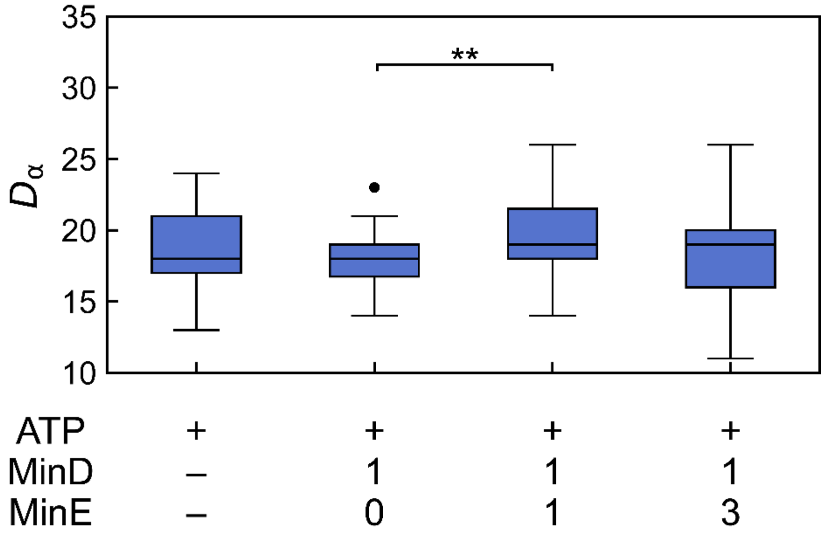


**Figure S9.** Apparent diffusion coefficient considering anomalous diffusion of TR-BSA with 2.5 mM ATP and different concentration of MinE and MinD proteins (see section S4 for the analysis). Min wave only exists under MinD: MinE = 1:1 [μM] (shown in red). Asterisks show significant differences in Welch's test (** $p < 0.01$) with false discovery rate (FDR) < 0.1 for multiple comparisons..

**Supporting Information Tables**

**Table S1.** TR-BSA diffusion under non-crowding conditions. See also Fig. 1.

| Solution condition | Bulk or droplet | Diffusion coefficient $D$ (Ave. ± S. D.) |
|---|---|---|
| 10 mg/mL BSA in PBS | bulk | 47.3 ± 5.7 $\mu m^2$/sec ($n$ = 10) |
| 10 mg/mL BSA in PBS | PC droplets | 49.3 ± 5.3 $\mu m^2$/sec ($n$ = 25) |

**Table S2.** TR-BSA diffusion under crowding conditions. See also Fig. 2 and S2.

| Solution condition | Bulk or droplet | Diffusion coefficient $D$ (Ave. ± S. D.) |
|---|---|---|
| 160 mg/mL BSA in PBS | bulk | $D$ = 25.3 ± 3.8 $\mu m^2$/sec ($n$ = 30) |
| 160 mg/mL BSA in TKM | bulk | $D$ = 22.5 ± 4.1 $\mu m^2$/sec ($n$ = 38) |

**Table S3.** Estimated $R^*$ and $D_{mini}$ for droplets with $R$ = 6 µm by the linear fitting of Fig. 2 and S2.

| Solution condition | Bulk or droplet | Diffusion parameter |
|---|---|---|
| PBS | PC droplets | $D_{mini}/D_{bulk}$ = 0.81, $R^*$ = 17.6 µm |
| TKM | PC droplets | $D_{mini}/D_{bulk}$ = 0.73, $R^*$ = 15.2 µm |
| PBS | E. coli polar lipids droplets | $D_{mini}/D_{bulk}$ = 0.75, $R^*$ = 20.0 µm |
| TKM | E. coli polar lipids droplets | $D_{mini}/D_{bulk}$ = 0.66, $R^*$ = 19.8 µm |
| PBS | PG droplets | $D_{mini}/D_{bulk}$ = 0.60, $R^*$ = 20.1 µm |
| TKM | PG droplets | $D_{mini}/D_{bulk}$ = 0.64, $R^*$ = 17.1 µm |

**Table S4.** Zeta potential of BSA and lipid membrane.

| Solution | PBS | TKM |
|---|---|---|
| 10 mg/mL BSA | −20.1 ± 0.7 mV (*n* = 5) | −17.4 ± 1.3 mV (*n* = 3) |
| 1 mg/mL PC vesicle | −7.0 ± 1.2 mV (*n* = 3) | −2.2 ± 0.7 mV (*n* = 3) |
| 1 mg/mL E. coli polar vesicle | −25.6 ± 0. 9 mV (*n* = 3) | −16.8 ± 0.1 mV (*n* = 3) |
| 1 mg/mL PG vesicle | −34.5 ± 0.9 mV (*n* = 3) | −22.7 ± 1.3 mV (*n* = 3) |

**Table S5.** Zeta potential of BSA solution with ATP.

| Solution | TKM |
|---|---|
| 10 mg/mL BSA in TKM | −17.4 ± 1.3 mV (*n* = 3) |
| 10 mg/mL BSA with ATP 250 µM in TKM | −17.8 ± 4.6 mV (*n* = 6) |
| 10 mg/mL BSA with ATP 2.5 mM in TKM | −16.6 ± 1.0 mV (*n* = 3) |

**Table S6.** TR-BSA diffusion under crowding conditions coexistence with Min system. Each solution dissolved in TKM. See Fig. 4 and 5.

| Solution condition | ATP 2.5 mM | MinD (µM) | MinE (µM) | Diffusion coefficient *D* (Ave. ± S.D.) |
|---|---|---|---|---|
| Control | − | − | − | 21.0 ± 2.7 µm$^2$/sec (*n* = 28) |
| MinD : MinE = 0:0 | + | − | − | 17.2 ± 1.8 µm$^2$/sec (*n* =36) |
| MinD : MinE = 1:0 | + | 1 | 0 | 17.1 ± 1.8 µm$^2$/sec (*n* = 36) |
| MinD : MinE = 1:1 | + | 1 | 1 | 18.4 ± 2.6 µm$^2$/sec (*n* = 35) |
| MinD : MinE = 1:3 | + | 1 | 3 | 17.0 ± 2.4 µm$^2$/sec (*n* = 36) |

**Table S7.** Results of the F-test for each combination of solution conditions under crowding conditions coexistence with Min system (** $p < 0.01$). See Fig.5

| Combination of solution condition | Significance | *p* value |
|---|---|---|
| MinD : MinE = 0:0 | | |
| MinD : MinE = 1:0 | n.s. | 0.4673 |
| MinD : MinE = 1:1 | * | 0.0206 |
| MinD : MinE = 1:3 | * | 0.0440 |
| MinD : MinE = 1:0 | | |
| MinD : MinE = 1:1 | * | 0.0170 |
| MinD : MinE = 1:3 | * | 0.0370 |
| MinD : MinE = 1:1 | | |
| MinD : MinE = 1:3 | n.s | 0.3621 |
